# Metformin enhances differentiation and function of skeletal muscle in models of Facioscapulohumeral Muscular Dystrophy (FSHD)

**DOI:** 10.64898/2026.07.30.736088

**Authors:** Joshua Greig, Jiani Qian, Philipp Heher, Peter S. Zammit

## Abstract

Facioscapulohumeral muscular dystrophy (FSHD) is one of the most prevalent inherited muscular dystrophies, for which there are no disease-modifying therapies. Metabolic perturbation, mitochondrial dysfunction, and oxidative stress are key contributors to FSHD pathology.

Here, the effects of the metabolic regulator and anti-diabetic drug Metformin on myogenesis and muscle function in human and murine models of FSHD were investigated. Metformin did not affect the proliferation rate of human control or patient-derived FSHD myoblasts but promoted their myogenic differentiation, increasing myotube formation and maturation. Metformin also enhanced the metabolic health and viability of myotubes. Mechanistic interrogation revealed reduced levels of mitochondrial reactive oxygen species and modified mitochondrial turnover. These cellular investigations were complemented with *in vivo* functional assessment in a murine model of FSHD, in which Metformin treated mice exhibited significantly improved muscle strength.

Collectively, these findings identify metabolic regulation as a therapeutically tractable feature of FSHD and demonstrate that Metformin improves muscle function in multiple models of FSHD via reduction of oxidative stress and augmentation of cellular metabolic fitness. These results provide insight into the therapeutic actions of Metformin and pre-clinical data to support its testing for repurposing in FSHD.

## Introduction

Facioscapulohumeral muscular dystrophy (FSHD) is the third most common inherited muscular dystrophy and is characterised by descending progressive skeletal muscle weakness and atrophy^1^. FSHD usually first presents in facial muscles and proceeds asymmetrically to muscles of the upper limbs and thorax^2–4^. The genetic aetiology of the condition is ectopic re-expression of the pioneer transcription factor *Double Homeobox 4 (DUX4*), which functions during zygotic genome activation ^4,5^. This aberrant re-expression of *DUX4* predominately originates from truncation of the D4Z4 microsatellite array located on chromosome 4, where each D4Z4 unit contains a DUX4 ORF^6^, combined with a permissive haplotype in *cis*, supplying a polyA signal^4,7,8^. Although *DUX4* is tightly restricted during postnatal development^8,9^, ectopic expression from the distal-most D4Z4 unit in skeletal muscle initiates a pathogenic transcriptional programme^4,7,8,10^. DUX4 is highly cytotoxic in somatic cells^8,11^ and induces a myriad of changes in fundamental cellular behaviours^7,12–16^. Despite major advances in understanding disease pathomechanisms, and potential therapies in clinical trial, there are no approved disease-modifying therapies for FSHD.

One key aspect of the DUX4 induced cellular pathology uncovered by recent work is the role of metabolic perturbations and mitochondrial dysfunction in FSHD pathology^7,13,16^. Mitochondria of FSHD skeletal muscle exhibit aberrantly high levels of reactive oxygen species (ROS) production, with reduced oxidative phosphorylation (OXPHOS) and ATP generation being a hallmark of FSHD myogenic cells ^8,13,16^. This defective mitochondria OXPHOS and elevated ROS result in elevated apoptosis in FSHD muscle cells and are likely central to the cellular pathology and atrophy of skeletal muscle in FSHD patients^13^. Importantly, mitochondrial perturbation may occur early during disease progression and contribute directly to DUX4-mediated cytotoxicity, rather than simply arising secondary to muscle degeneration^13,16^. This concept is supported by studies demonstrating that modulation of mitochondrial activity or oxidative stress can partially rescue DUX4-associated phenotypes in experimental systems^13,17,18^. Collectively, these findings identify metabolic dysfunction as a potential target in FSHD muscle. Indeed, mitochondrial dysfunction and general metabolic dystasis are documented in other muscular dystrophies including Duchenne (DMD) and Beckers muscular dystrophy ^19–25^ This suggests that general metabolic dysfunction may be a key feature of muscular dystrophies and thus metabolic regulators may have general utility as adjunctive theraputics^24^.

Such a metabolic regulator is Metformin, a biguanide compound derived from goats’ rue (French lilac) which has been known to have anti-diabetic effects since ancient times^26^. It is the first front-line pharmacological treatment prescribed for Type 2 Diabetes Mellitus ^27^ since first being clinically and scientifically described in 1957^26^. In Diabetes, Metformin bypasses the defective insulin receptor signalling pathway to mobilise the glucose uptake transporter GLUT4 to the plasma membrane, increasing uptake of glucose from the bloodstream to alleviate hyperglycaemia^27,28^. Increasingly, Metformin is being recognised to exert broader effects on cellular metabolism and stress adaptation beyond enhancing glucose uptake ^29–35^. Metformin acts on the energy sensor AMP-activated protein Kinase (AMPK) which detects the ATP:AMP ratio and thereby acts either directly or indirectly on the complexes of the electron transport chain in mitochondria^27,31,32,35–40^. There is also evidence that Metformin can regulate oxidative stress, mitochondrial quality control, autophagy and inflammatory signalling pathways in multiple tissues^37,41,42^. Additionally, alterations to metabolic activity in hepatocytes may be beneficial for organismal health^43^.

Metformin has been tested in other muscular dystrophies, notably DMD, with mixed results. In some mouse studies, Metformin has been shown to be beneficial in the *mdx* model of DMD, with enhancement of skeletal muscle function^29,44,45^. However, other studies in both mouse and human patients report that either Metformin has no detectable effect or is detrimental to muscle hypertrophy following a strength training regime^46,47^. What is evident is that a range of Metformin dosage regimes have been used, indicating that there is scope for optimisation.

Here, we examined the effectiveness of Metformin in FSHD, in both *in vitro* human patient-derived myotubes and *in vivo* using the *FLExDUX4* mouse model of FSHD to assess muscle function. First, an empirical measurement of the dosage effect relationship of Metformin on myogenesis was determined in human myoblasts. Metformin exhibited a bi-phasic hormetic dosage response curve, with higher dosages being detrimental to differentiation of both human control and FSHD patient-derived myoblasts but critically, not due to cyto-toxicity. At lower dosages, beneficial effects of Metformin on myogenesis occurred over a range of dosages below a key switch threshold, indicating a wide therapeutic window. Metformin did not adversely affect myoblast proliferation, and at the highest myogenically beneficial dosage, enhanced metabolic activity. Investigation into the underlying mechanism indicated that reduced mitochondrial ROS production and enhanced mitophagy were key to Metformin’s effect. Crucially, our *in vitro* analysis was complemented with *in vivo* data demonstrating that Metformin increased the strength of skeletal muscle in the *FLExDUX4* FSHD mouse model. Together our data demonstrate the effectiveness of Metformin in addressing the impaired myogenesis in FSHD across model systems, and this work acts as a pre-clinical case for exploring Metformin for drug reproposing for FSHD^48^.

## Results

### Metformin exhibits a bi-phasic hormetic dosage dependent effect on myogenic differentiation

We first determined the efficacy of a large range of Metformin dosages empirically. A 96-well plate-reader based assay was developed to assess myogenesis (Fig. 1). Each myoblast cell line was seeded with a comparable control or FSHD in triplicate rows of ten wells (Fig. 1A). To provide a ceiling to the dosage, and thus an indication of the operational and effective therapeutic window (optimal range), a 10x fold dilution interval was used, starting from the highest commonly reported dosage^49^. Each dosage of Metformin was administered with differentiation medium (timepoint 0 h) for the duration of differentiation (72 h total). Differentiation of myotubes was then assessed by immunolabelling for Myosin heavy chain (MyHC) and nuclei stained with Hoechst and a plate reader used to measure whole-well signal of both MyHC and myonuclei for normalization (MyHC/nuclei) then expressed relative to the vehicle-only control.

**Fig 1:**
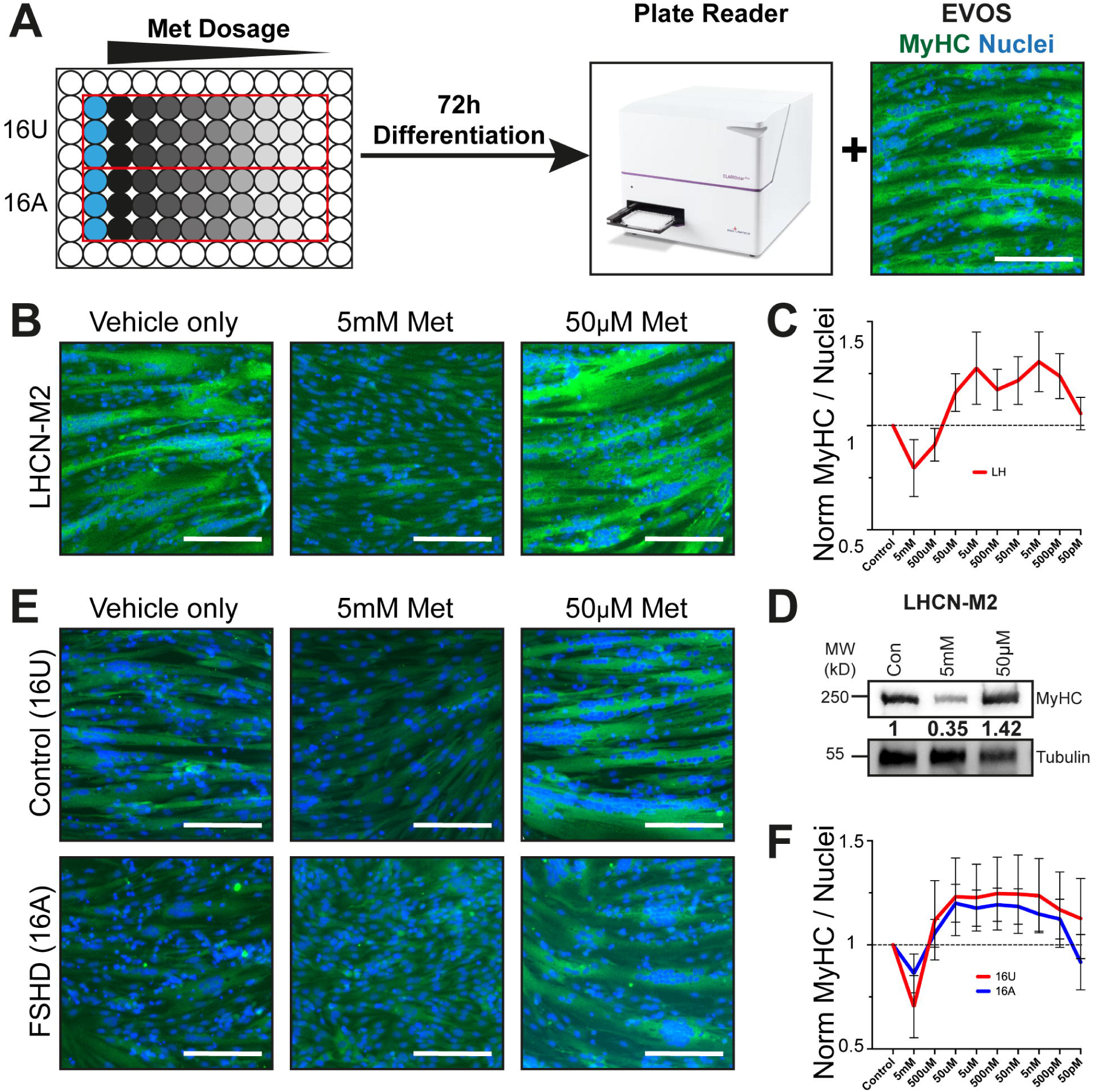
Metformin dosage effects myogenesis in both control and FSHD myotubes. (A) Illustration of the 96-well plate based myogenesis assay, each plate contains the FSHD immortalised myoblast cell line (16A) and its respective control (16U). Vehicle-only (Blue Wells, left) and each dosage (reducing across the plate left to right) are administered to three separate wells constituting the technical replicates for each biological replicate. At the end point, cells are fixed, immunolabelled and imaged on both a plate reader and an Invitrogen EVOS™ M5000 Imaging platform (EVOS). (B) Representative merged images of LHCN-M2 myotubes differentiated in the presence of Vehicle only (Water), 5 mM or 50 μM Metformin, immunolabelled for MyHC (Green) and nuclei stained with Hoechst (Blue). Scale bars represent 250 µm. (C) Quantification of the differentiation of LHCN-M2 myotubes at the different Metformin dosages. Values adjusted for cell number using nuclei staining (MyHC/Hoechst) and normalized to the differentiation of Vehicle-only control. Each point represents the normalized mean value for the indicated dosages across all experiments with bars representing ±SEM. The dotted line represents the differentiation level in control vehicle-only (n = 5 biological replicates). (D) Representative immunoblot for MyHC and Actin of control myotubes (LHCN-M2) differentiated in the presence of Vehicle only (Con), 5 mM or 50 μM Metformin. Normalized mean optical density (OD) values for MyHC are given below (n = 3 biological replicates). The migration positions of molecular weight (MW) markers are indicated at the left in kilodaltons (kD). (E) Representative merged images of control (16U) and FSHD (16A) myotubes differentiated in the presence of Vehicle only (Water) or 5 mM or 50 μM Metformin, immunolabeled for MyHC and nuclei stained with Hoechst (Blue). Scale bars represent 250 µm. (F) Quantification of the differentiation of FSHD myotubes at the different Metformin dosages. Values adjusted for cell number using nuclei staining (MyHC/Hoechst) and normalized to the differentiation of Vehicle-only control. Each point represents the normalized mean value for the indicated dosages across all experiments with bars representing ±SEM. The dotted line represents the differentiation level in control vehicle-only (n = 6 biological replicates).

The human control myoblast cell line LH (LHCN-M2) was first used to assess the efficacy of Metformin (Fig. 1B). Myoblasts exposed to the highest dosage of Metformin (5 mM) exhibited a distinct lack of large myotubes and low expression of MyHC (Fig. 1B, middle). Critically, nuclei were still present, and the colour change of the cell medium indicated metabolic activity, suggesting higher dosages of Metformin were not overtly cytotoxic. While higher dosage of Metformin (5 mM - 500 μM) were detrimental to myogenesis, lower dosages were beneficial (Fig. 1B, middle). This effect persisted through 10x fold magnitude dilutions down to 50 pM, at which dosage differentiation was indistinguishable from control vehicle-only (Fig. 1C). This dosage response pattern is consistent with a bi-phasic or hormetic drug effect^50^. Western blot assay confirmed the effect of Metformin on MyHC (Fig. 1D), revealing that higher dosages reduced MyHC levels by ∼65% (optical density measurement of 0.35 relative to control) while the highest beneficial dosages tested (50 μM) resulted in a ∼42% increase (1.42 relative to control) (Fig. 1D).

To investigate the efficacy of Metformin in FSHD, a patient-derived immortalized myoblast line with a contracted D4Z4 allele (16A) and a sibling control (16U)^51^ were subject to the myogenesis assay (Fig. 1E). The FSHD 16A myoblasts yielded hypotrophic myotubes compared with the 16U in the vehicle-only control, as previously observed^13^.

The higher Metformin dosages yielded a significant reduction in morphologically distinct myotubes and reduced expression of MyHC. Conversely, lower dosages enhanced myogenesis in both the control (16U) and FSHD affected (16A) myoblast line (Fig. 1E). Normalization was performed within each myoblast line to its respective vehicle-only dosage control thus permitting comparison between the lines for dosage response rather than innate differences in differentiation capacity. Higher dosages were again detrimental, with lower dosages being beneficial over a wide magnitude of dosages before being indistinguishable from control vehicle-only at the lowest dosage (Fig. 1F).

In summary, Metformin has a hormetic dosage dependent effect on myogenesis, and lower dosages are beneficial for both human control and FSHD myotubes.

### Metformin does not perturb myoblast proliferation and enhances cell viability

As Metformin is administered continuously and systemically, consideration was required for the effects of the compound beyond myogenesis. We thus focused three fundamental cellular processes: proliferation, apoptosis, and general cellular metabolic health.

FSHD (16A) and sibling matched control (16U) myoblasts were subjected to Metformin dosages for 24 h and the myoblast proliferation rate measured by EdU incorporation (Fig. 2A). Two dosages were used: the highest dosage from the myogenesis assay (5 mM) or the highest that yielded a beneficial effect on myogenesis (50 μM). The percentage of total nuclei containing EdU was calculated and neither dosage in either myoblast cell line perturbed EdU incorporation (Fig. 2B).

**Fig 2:**
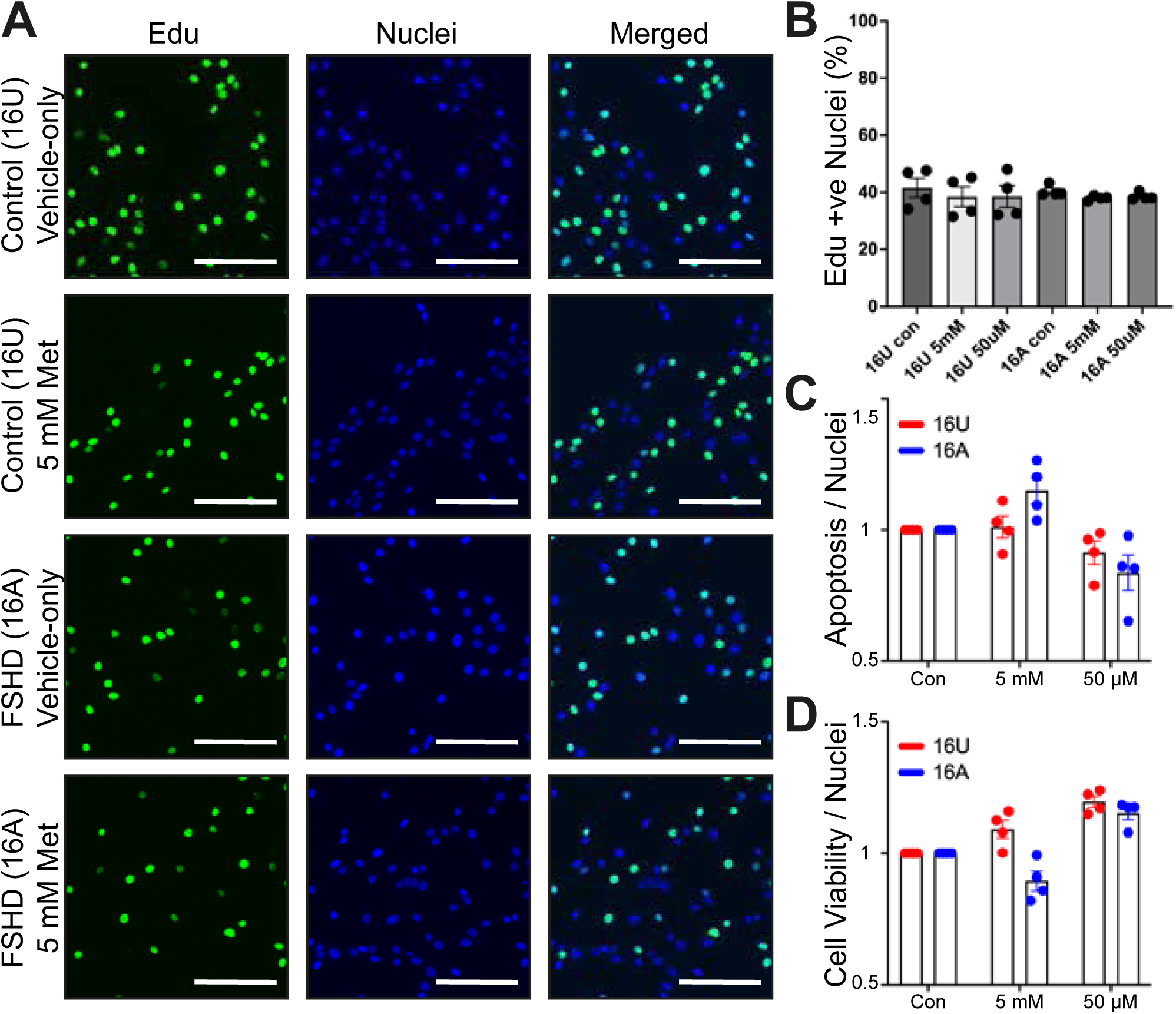
Metformin does not perturb myoblast proliferation and enhances myotube viability. (A) Representative images of control (16U) and FSHD (16A) myoblasts in standard growth medium with vehicle (Vehicle-only) or 5 mM of Metformin (5 mM Met) for 24 h and pulsed with EdU for 2 h prior to fixation. EdU incorporated nuclei (Green) and total nuclei stained with Hoechst (Blue) with the frequency indicated in the merged image. Scale bars represent 250 µm. (B) Quantification of nuclei containing EdU (EdU +ve) as a percentage of total myoblast nuclei (%). Each point represents the mean value from a biological replicate for each myoblast line in each condition with the bars representing ±SEM (n = 4 biological replicates). Statistical analysis was performed using a one-way ANOVA with a Tukey post-hoc test. (C) Quantification of Apoptosis in control (16U, red) and FSHD (16A, blue) myotubes at the endpoint of differentiation (72 h) measured using the Annexin V Apoptosis luminescent assay. Myotubes differentiated with either vehicle-only (Con) or with 5 mM or 50 μM Metformin. Whole-well Luminescent signal values adjusted for cell number using Whole-well Hoechst nuclei staining (Apoptosis/Nuclei) and each Metformin dosage normalised to vehicle-only (Con) within each myotube cell line. Each point represents the normalized mean value from a biological replicate with the bars representing ±SEM (n = 4 biological replicates). (D) Quantification of Cell Viability in control (16U, red) and FSHD (16A, blue) myotubes at the endpoint of differentiation (72 h) measured using the RealTime-Glo luminescent assay. Myotubes differentiated with either vehicle-only (Con) or with 5 mM or 50 μM Metformin. Whole-well Luminescent signal values adjusted for cell number using whole-well Hoechst nuclei staining (Cell Viability/Nuclei) and each Metformin dosage normalised to vehicle-only (Con) within each myotube cell line. Each point represents the normalized mean value from a biological replicate with the bars representing ±SEM (n = 4 biological replicates).

To assess the effect of Metformin on apoptosis, the Annexin V based luminescence assay normalized for nuclei was used with the two dosages of 5 mM and 50 μM (Fig. 2C). A mild increase in apoptosis was detected at the 5 mM dosage of Metformin, relative to the vehicle-only control (con), which was more apparent in the FSHD 16A myotubes (Fig. 2C). However, a reduction in apoptosis was observed in both FSHD (16A) and control (16U) myotubes at the beneficial dosage (50 μM) (Fig. 2C).

To complement these analyses, a general assessment of cellular metabolic health was undertaken using a cell viability assay that measures the reducing capacity of the cell as a proxy for metabolic health/activity (Fig. 2D). At 5 mM Metformin a decrease of cell viability was detected in the FSHD myotubes (mean value of 0.89, SD 0.068, relative to vehicle-only control) (Fig. 2D). However, both the FSHD (16A) and control (16U) myotubes exhibited an increase in cell viability at 50 μM (mean value of 1.17, SD 0.045, relative to control vehicle only) (Fig. 2D).

Overall, Metformin at the myogenically beneficial 50 μM dosage had no adverse effects on the proliferation rate, and a beneficial effect on cell viability and apoptosis.

### Metformin reduces mitochondrial ROS and modulates mitophagy

As Metformin is purported to also act via the mitochondria, the next step was to examine mitochondrial activity as a potential explanatory mechanism. To assess activity and function of mitochondria, membrane potential (ΔΨm), total intracellular Reactive Oxygen Species (ROS) and mitochondrial-specific ROS (mitoROS) were assayed at the 72 h endpoint of differentiation^13^ (Fig. 3).

**Fig 3:**
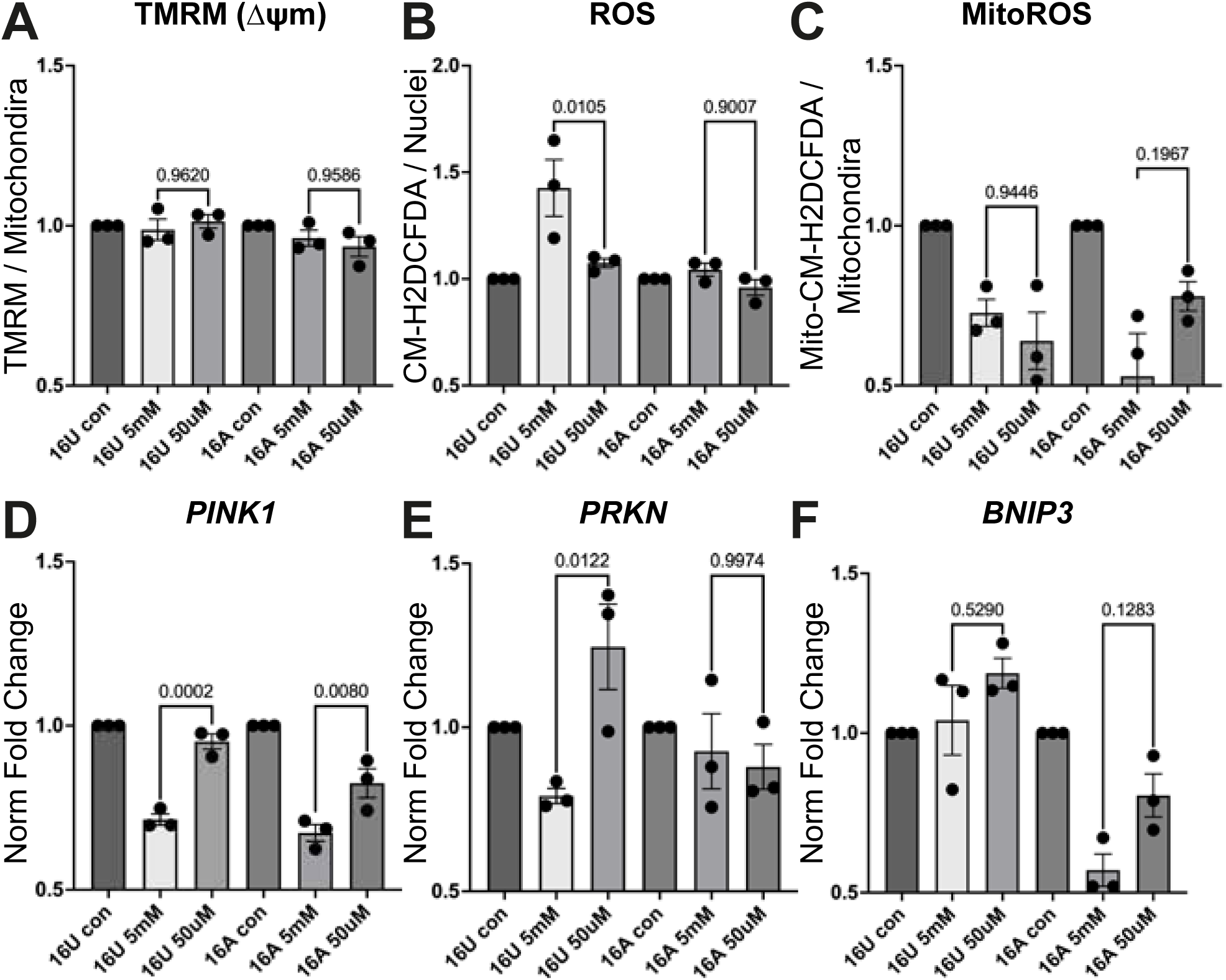
Mitochondrial ROS production is reduced and mitophagy stimulated by Metformin. (A-C) Control (16U) and FSHD (16A) myotubes differentiated with either vehicle-only (Con), 5 mM or 50 μM Metformin at the endpoint of differentiation (72 h) assayed for (**A**) Mitochondrial membrane potential (Δψm) using tetramethylrhodamine methyl ester (TMRM) fluorescence, (**B**) total Reactive Oxygen Species (ROS) using CM-H_2_DCFDA fluorescence or (**C**) Mitochondrial Reactive Oxygen Species using MitoTracker® Red CM-H_2_XROS (Mito-CM-H_2_DCFDA) fluorescence. Whole-well fluorescent signal values adjusted for cell number using whole-well Hoechst nuclei staining and secondly normalised for total mitochondrial content using MitoTracker Deep Red. Each Metformin dosage normalised to vehicle-only (con) within each myotube line. Each point represents the normalized mean value from a biological replicate with the bars representing ±SEM (n = 4 biological replicates). Statistical analysis was performed using a one-way ANOVA with a Tukey post-hoc test, individual pairwise *p*-values from this post-hoc test are displayed. (**D-F**) Control (16U) and FSHD (16A) myotubes differentiated with either vehicle-only (Con), 5 mM or 50 μM Metformin at the endpoint of differentiation (72 h) assayed using RT-qPCR for the mitophagy markers (**D**) *PINK1* (**E**) *PRKN* (encodes Parkin) and (**F**) *BNIP3*. Values expressed as normalised transcript fold change for each Metformin dosage relative to vehicle-only (con) within each myotube cell line. Each point represents the normalized mean value from a biological replicate with the bars representing ±SEM. (n = 3 biological replicates). Statistical analysis was performed using a one-way ANOVA with a Tukey post-hoc test, individual pairwise *p*-values from this post-hoc test are displayed.

When membrane potential (ΔΨm) in control (16U) and FSHD (16A) myotubes was assayed with tetramethylrhodamine methyl ester (TMRM) fluorescence, no significant difference was detected between the highest dosage (5 mM) and the highest beneficial dosage (50 μM) for both myotubes cell lines (Fig. 3A).

Next, total cytoplasmic ROS was examined using the general ROS indicator CM-H2DCFDA (Fig. 3B). Curiously, the highest dosage of Metformin produced an elevation of ROS in the control (16U) relative to both control vehicle only (con) and the beneficial (50 μM) dosage (p = 0.01). This was not observed in FSHD (16A) myotubes which exhibited no differences in ROS between the Metformin dosages (Fig. 3B).

To examine ROS generated from mitochondria directly, the targeted probe MitoTracker® Red CM-H_2_XROS^13^ was used (Fig. 3C). This revealed a substantial and significant decrease in the mitoROS production in both control and FSHD myotubes with either dose of Metformin. At 50 μM Metformin, the mitoROS levels were reduced by over ∼36% (0.64 SD 0.15, normalised to respective vehicle-only control) for the control 16U myotubes and by ∼23% (0.77 SD 0.078, normalised to respective vehicle-only control,) for the FSHD myotubes (Fig. 3C). Together, these data indicate that Metformin does not alter mitochondria membrane potential, a reliable proxy for function, but does reduce ROS generated by the electron transport chain (ETC).

To investigate effects that Metformin has upon mitochondrial turnover, expression of mitophagy/autophagy genes was examined at the endpoint of differentiation (Fig. 3D-E). PINK1/Parkin are the classical canonical pathway that regulates the mitophagy turnover of mitochondria^52,53^. Measurement of *PINK1/PRKN* (the latter encoding Parkin) expression under the different Metformin dosages revealed a significant reduction in expression which was correlated with Metformin dosage in both FSHD and respective control myotubes (Fig. 3D). An inverse relationship between the highest and lower Metformin dosages was detected in control (16U) myotubes. The Contrasts between *PINK1* and *PRKN* with the higher 5 mM dosage suggests that higher levels of Metformin may be saturating the mitophagy pathway (Fig. 3E). Curiously, FSHD myotubes (16A) did not exhibit the same dosage relationship pattern with *PRKN* as with *PINK1* (Fig. 3D+E) suggestive that FSHD myotubes may have a discord with the coupling of PINK1 and Parkin.

To explore activity of the mitophagy pathway that operates in parallel but separate from the PINK1/Parkin pathway, the expression of the non-canonical mitophagy and hypoxia responsive adaptor BNIP3 was examined^54^ (Fig. 3F). In control (16U) myotubes, a mild elevation of *BNIP3* was detected at the lower Metformin dosage but no significant change at the higher. By contrast a significant reduction of *BNIP3* was detected in the FSHD (16A) myotubes at both Metformin dosages (Fig. 3F). This suggests that FSHD myotubes may be utilizing a different parallel pathway for mitophagy relative to the control myotubes.

Together, these investigations show that Metformin has a limited direct effect upon mitochondrial function, indicated by ΔΨm measured with TMRM, but a significant effect upon reducing the mitochondrial specific ROS production. Moreover, changes in expression of canonical *PINK1/PRKN* and non-canonical *BNIP3* are indicative of involvement of mitochondrial turnover by mitophagy, albeit with some potential difference in pathway choice between FSHD and control myotubes.

### Metformin enhances muscle strength in a mouse model of FSHD

To investigate the prospective efficacy and effectiveness of Metformin in treatment of FSHD pathology, the effect on skeletal muscle function in a murine model of FSHD *in vivo* was performed. To calculate a dosage for the mouse study from the dose used in the *in vitro* human myoblast data required consideration of bioavailability and drug metabolism. The highest dosage that produced a beneficial effect *in vitro* was 50 μM (Fig. 1), and so a dosage of 100 mg/kg/day (100 mkd) was calculated. This was administered by oral gavage to replicate the oral route used for Metformin in patients with Type 2 Diabetes. Adult *FLExDUX4.CRE* mice (12-14 weeks of age) were treated with either Vehicle (n = 10 animals) or 100 mg/kg Metformin (n = 10 animals) via daily oral gavage for 28 days without induction of DUX4^55^, and compared to vehicle-only treated wild-type CRE (WT.CRE) mice (n = 10 animals) (Fig. 4). Muscle function was assessed by measuring endurance in treadmill exhaustion and strength by isometric force generation in the hindlimb under electrical stimulation to measure the contractile force of the plantarflexor muscle group (Fig. 4A).

**Fig 4:**
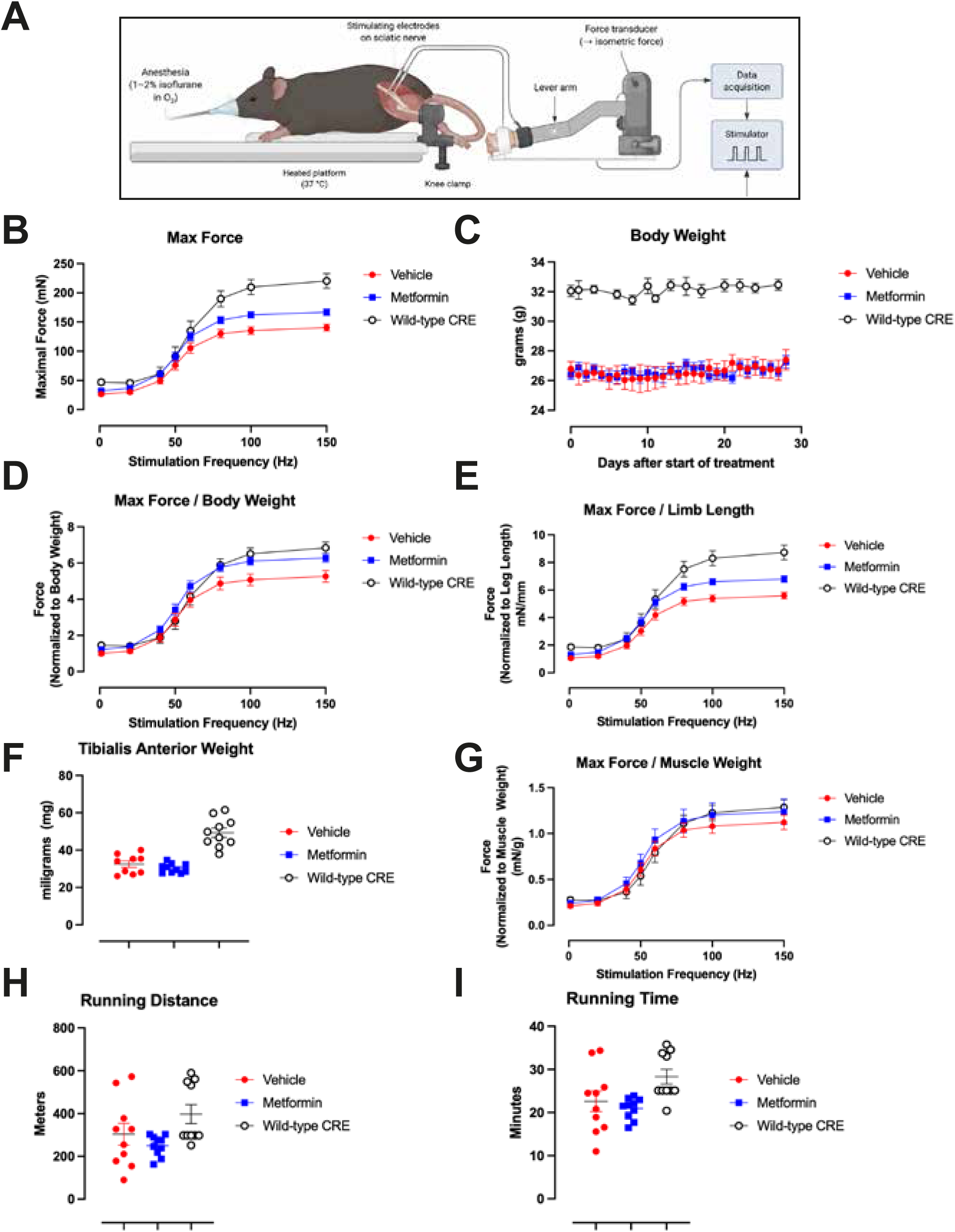
Metformin administration improves muscle force in a mouse model of FSHD. (A) Schematic of the *in vivo* muscle function assay Adult (12-14 weeks of age) *FLExDUX4.CRE* mice were treated with either Vehicle (red squares, n = 10 animals), or 100 mg/kg Metformin (Blue squares, n = 10 animals) via daily oral gavage for 28 days and compared to untreated wild-type CRE (WT.CRE) littermates (open circles, n = 10 animals). (B) Mean maximal contractile force (mN) generated in the *in vivo* muscle function assay relative to the stimulation frequency (Hz) of the hindleg of each mouse. Each point represents the mean value from all animals used in each condition with bars representing ±SEM. (C) Mean body weight of each mouse in each condition on each day during the 28-day time-course of the study. Each point represents the mean ±SEM value. (D) Mean hindlimb contractile force normalized to the body weight of each mouse in each condition. (E) Mean hindlimb contractile force normalized to the length of the hindleg in each mouse in each condition. (F) Tibialis Anterior muscle weight of each mouse in each condition measured post dissection at the endpoint of the study. Each point represents the measurement of each muscle, where the black line represents the mean ±SEM. (G) Mean hindlimb contractile force normalized to the Tibialis Anterior muscle weight of each mouse in each condition. (H) Running distance of each mouse on the treadmill assay in each condition measured at the endpoint of the study. Each point represents the measurement from each animal in each condition with the line representing the mean ±SEM. (I) Running time of each mouse on the treadmill assay in each condition measured at the endpoint of the study. Each point represents the measurement from each animal in each condition with the line representing the mean ±SEM.

Isometric hindlimb contractile force measured at the end point revealed a significant difference between the Metformin treated and vehicle treaded *FLExDUX4* mice in the total force generated across the range of stimulation frequencies (Fig. 4B). However, this did not reach the equivalent force generated by the untreated WT strain control. *FLExDUX4* mice were smaller in size and mass than their WT littermates (Fig. 4C). When force measurement was corrected for body mass, the Metformin treated *FLExDUX4* mice generated a force significantly greater than the vehicle-only control (Fig. 4D), which was also indistinguishable from the normalized values for the WT.CRE external control (Fig. 4D).

To further analyse these force-frequency relationships, a sigmoidal nonlinear regression model (4 parameter logistic) was used with comparison of the curves by extra sum-of-squares F test ^56^. When the WT.CRE and *FLExDUX4*-Vehicle, or *FLExDUX4*-Vehicle and *FLExDUX4*-Metformin curves were compared both pairs were significantly different (p < 0.0001 for both) indicating that the datasets were not described by a single curve response. However, the response curves were no longer different between the WT.CRE and *FLExDUX4*-Metformin treated mice (p = 0.15). That the WT.CRE and *FLExDUX4*-Metformin fit a single response curve indicates that Metformin restores the force-frequency response observed in the WT.CRE animals when data is corrected for total body weight (Fig. 4D).

To further explore comparative correctives, total force was normalized to the length of the hindlimb (Fig 4E) Normalized force generation of the Metformin treated *FLExDUX4* mice was significantly greater than the vehicle-only *FLExDUX4*, and tending towards, values in the untreated WT.CRE external control (Fig. 4E).

The weight of the tibialis anterior muscle measured post-mortem was lower in the *FLExDUX4* stain compared to the WT.CRE littermates (Fig. 4F). When normalised to muscle weight, force generation of the Metformin treated *FLExDUX4* mice and vehicle-only *FLExDUX4*, was comparable to values in the untreated WT.CRE external control (Fig. 4G). However, the tibialis anterior is only one of the several muscles of the plantarflexor muscle group which was measured in this isometric hindlimb contractile force assay.

Measurement of functional endurance in the mice was undertaken with a treadmill exhaustion assay. Metformin treated *FLExDUX4* mice did not outperform the Vehicle-only control mice in either running distance (Fig. 4H) nor running time to exhaustion (Fig. 4I).

Overall, this *in vivo* functional testing showed that Metformin treatment increased skeletal muscle strength in a murine model of FSHD.

## Discussion

In this study, we found that Metformin exerts a bi-phasic hormetic effect on the myogenesis of human myoblasts and beneficial dosages are in evidence across a magnitude fold range. Metformin also had no adverse effects on proliferation and a beneficial effect on cell viability. Mechanistic interrogations indicated reduced mitochondrial ROS and apparent altered mitophagy in human myotubes. Critically, these findings are supported by *in vivo* examination in the *FLExDUX4* mouse model of FSHD that showed a significant increase in the contractile force generated by mice treated with Metformin. Together, our data provide evidence for further investigation and a pre-clinical rationale for exploration for the rapid testing of Metformin for drug repurposing in FSHD patients.

While this is the first investigation in FSHD, Metformin has been tested in DMD with conflicting outcomes: reporting beneficial, ineffective, or detrimental effects^44,46,47,57^. While the aetiology of FSHD and DMD differs, it is important to note that these studies also used different dosage regimens of Metformin. Given our observations, it is likely that the differing results are a consequence of Metformin exhibiting a hormetic drug dose response curve (Fig. 1). Indeed, a previous report identified significant dose effects of Metformin in murine-derived myoblasts ^49^ which tallies with the broader dosage range findings in human myotubes here. Moreover, this hormetic dosage response effect provides an explanation for the blunted muscle hypertrophy observed in older human patients who underwent resistance training with a Metformin dosage of 1700 mg/day^47^. Patients with type 2 diabetes are reported to be at a higher risk from muscle loss and weakness^58^, with many clinical investigations now classifying the effect of diabetes on skeletal muscle as a distinct diabetic myopathy condition. Since most diabetic patients will be on high doses of metformin, it is intriguing to speculate as to whether this may be a contributory factor given the detrimental effects on human myogenesis even observed in myoblasts from healthy individuals. Patients with both Type 2 Diabetes and FSHD may need to reduce their Metformin dose to achieve a compromise for treating both Type 2 Diabetes and supporting muscle in FSHD.

Type 2 Diabetic patients typically start at a dosage of 500 mg per day and ramp up to 2000-2500 mg per day ^35^. Based upon a calculation of the human equivalent dosage (HED)^59^ from our mouse study, this would exceed the myogenically beneficial dosage threshold identified here. Given the 10x fold dilutions performed for the initial myogenesis assay and the enhanced myogenesis detected across a range of dosages below 50 μM, low Metformin dosing seems more appropriate to protect skeletal muscle. We selected the highest beneficial dosage from the *in vitro* experiments to inform the dosage selected for the mouse study. We reasoned that even if more Metformin failed to be absorbed or was excreted, thus reducing its bioavailability, the reduced dosage would still yield a beneficial effect. Metformin is being explored as a drug repurposing candidate for DMD (e.g. by Duchenne muscular dystrophy UK) and our work indicates that doses equivalent to those used for Type 2 Diabetes may be too high when testing to repurpose this inexpensive and readily available compound.

The mechanisms of action of Metformin have been subject of debate and investigation since it was fist administered clinically some 70 years ago ^26,27^. The prevailing view is that it acts as a cellular energy regulator by stimulating the activity of AMP kinase (AMPK) leading to elevation of glucose uptake and metabolic activity^36,39,41,60^. However, there is no indication that this action upon AMPK is direct nor that it is the sole mechanism^32,35,39–41^. Metformin is also posited to directly and irreversibly inhibit Complex-I of the electron transport chain (ETC)^35,39,60^, although some of these early studies primarily used isolated and purified mitochondria with millimolar concentration of Metformin^61^. However, there are compelling recent studies indicating that inhibition of Complex I may be one of the primary mechanisms of action^62^. Nonetheless, if Metformin is exhibiting a low affinity inhibitory effect on Complex I, it would amount to the same overall cellular outcome, since AMPK activity is sensitive to the ETC and particularly Complex-I activity^63^. Our findings support Metformin acting upon Complex I as reduced mitoROS was detected (See Fig. 3) and Complex-I is one of the main sources of ROS production in mitochondria^64^. Identifying that a myogenically beneficial dosage (50 μM) of Metformin also enhanced cell viability (See Fig. 2) further supports metabolic rewiring or metabolic shifting. Aberrant ROS production via OXPHOS is central to pathology in FSHD^13,17^, and so a compound like Metformin would have a beneficial role in such a condition.

A key finding of this study is that Metformin is beneficial when administered systemically, translating our *in vitro* data to the *in vivo* mouse system (See Fig. 4). Previous mouse studies have used a range of Metformin dosages from 50-500 mg/kg/day with variable outcomes^65,66^. For this study, calculation of dosage was based upon the *in vitro* data, with additional consideration of bioavailability based upon the oral gavage route of administration. This resulted in an improvement in muscle strength but not endurance of *FLExDUX4* mice (Fig. 4). This could indicate muscle fibre type specific effects, with Metformin having a beneficial effect on the fast-twitch (Type II) fibres that generate more contractile force^67^, rather than the slow-twitch (Type I). Metformin preferentially affects activity of fast-twitch fibres, which are more reliant on glycolysis resulting from higher AMPK activity^68^. A non-mutually exclusive explanation is that Metformin is also preserving muscle by limiting atrophy.

Metformin alone did not affect the hypotrophic phenotype of FSHD as no gross increase in muscle mass was observed in the treated mice (Fig. 4). FSHD patients have a fast-twitch fibre defect, with FSHD muscle favouring slow-twitch fibres^69^. One intriguing prospect would be to combine an optimal dose of Metformin with strength/resistance training regimes to enhance muscle function in patients, or at the very least, to abate the rate of atrophy to preserve function.

In conclusion, use of the anti-diabetic compound Metformin improved phenotypes associated with FSHD *in vitro* and *in vivo*. Metformin has a hormetic dosage response relationship in both healthy and FSHD myotubes. Further, this effect occurs across a range of dosages below 50 μM indicating a wide therapeutic window. This is coupled with no adverse effects on cellular proliferation and a beneficial effect on cellular metabolism. Importantly, Metformin increased muscle strength in a mouse model of FSHD. Since Metformin has been used clinically for treatment of Type 2 Diabetes for 70 years, the safety profile is well established. Thus, this study provides a strong pre-clinical case for the exploration of Metformin as an inexpensive and readily available drug for reproposing in FSHD patients.

## Materials and Methods

### Cell culture

The FSHD patient-derived cellular model was the sibling-matched immortalised model derived from biceps muscle, with D4Z4-contracted FSHD clone 16A and uncontracted control clone 16U^51^. The control healthy myoblast line was the LHCN-M2 line derived from pectoral muscle^70^.

Human myoblast lines were cultured in Skeletal Muscle Cell Growth Medium (Promocell, Heidelberg, Germany) supplemented with 10% foetal bovine serum (FBS; ThermoFisher Scientific, MA, USA), 50 μg/mL fetuin (bovine), 10 ng/mL epidermal growth factor (recombinant human), 1 ng/mL basic fibroblast growth factor (recombinant human), 10 μg/mL insulin (recombinant human), 0.4 μg/mL dexamethasone (all added as PromoCell Supplement Mix) and 50 μg/mL gentamycin (Sigma Aldrich) in a humidified incubator at 37°C with 5% CO_2_. Myoblast lines were kept subconfluent in routine culture and passaged at maximum 70% confluency. To induce differentiation, myoblasts seeded for full confluency were washed twice with phosphate buffered saline (PBS) and placed in Dulbecco’s Modified Eagle Medium (DMEM) GlutaMax (ThermoFisher Scientific) supplemented with 10 μg/ml recombinant human insulin (Sigma-Aldrich).

### Animal Welfare

The study was conducted by Myologica (Myologica LLC). Sykesville, MD, USA. Myologica conducts laboratory animal research within Noble Life Science, who have aa PHS approved Animal Welfare Assurance issued by the Office of Laboratory Animal Welfare (OLAW) and is accredited by AAALAC. Noble Life Science in Sykesville, MD maintains a formal Institutional Animal Care and Use Committee (I.A.C.U.C.) that meets to review proposed studies, as well as current and ongoing studies in their laboratories.

This study complied with all appropriate parts of the Animal Welfare Act of 1966 (P.L. 89-544), as amended by the Animal Welfare Act of 1970 (P.L. 91-579) and the 1976 amendments to the Animal Welfare Act (P.L. 94-279) - 9 CFR Chapter I (January 1, 1998 edition) as well as the International Guiding Principles for Biomedical Research Involving Animals developed by the Council for International Organizations of Medical Sciences.

### Study Design

Male *FLExDUX4.CRE* mice were be assigned to one of two groups based on body weight. Mice were treated with vehicle (H_2_O) or Metformin alone (PO daily by oral gavage). Tamoxifen was not administered. Additionally, one group of littermate CRE WT mice served as a reference and were treated with vehicle alone via daily oral gavage). After 4 weeks of treatment, mice underwent treadmill endurance testing, followed by muscle function testing and then euthanized.

Mice were weighed, then held via tail and scruff restraint. For oral gavage, a 22G flexible gavage needle with a soft foam tip was used to administer Metformin or vehicle at 10 ml/kg volume. The needle was inserted to the right or left side of the mouth, down the oesophagus and the entire volume discharged into the stomach. The animal was then returned to its cage.

### Treadmill Exhaustion Assessment

Running capacity was assessed using a treadmill run-to-exhaustion protocol. Mice were acclimated to the treadmill (LE8710, Panlabs, Spain) for 3 days prior to measurement. On the first day, mice were placed on the treadmill set at 0 m/min for 5 min at no inclination. The following two days, mice were placed on the treadmill set at 5 m/min for 5 min at no inclination. Following the acclimation period, mice were placed on the treadmill with the speed was set at 5 m/min for 5 min at no inclination, the speed was then increased to 25m/min and mice were run until exhaustion.

### In Vivo Muscle Function

Following isoflurane sedation, muscle performance was measured *in vivo* with a 305C muscle lever system (Aurora Scientific Inc., Aurora, CAN). Mice were anesthetized via isoflurane inhalation (∼3-4.5%, or to effect) and placed on a thermostatically controlled table where anaesthesia maintenance was via nosecone (∼1-3% isoflurane, or to effect). The right knee was isolated using a pin pressed against the tibial head and the foot firmly fixed to a footplate on the motor shaft. For the plantarflexor muscle group, contractions were elicited by percutaneous electrical stimulation of the sciatic nerve. Optimal isometric twitch torque was determined by increasing the current with a minimum of 30 s between each contraction to avoid fatigue. A series of stimulations were performed at increasing frequency of stimulation (0.2 ms pulse, 500 ms train duration): 1, 20, 40, 50, 60, 80, 100, 150 Hz. Maximal peak isometric force at each frequency, and force-frequency relationship were plotted.

### Antibodies and Nuclear Stain

Primary antibodies were mouse anti-MyHC (MF-20; Developmental Studies Hybridoma Bank, IA, USA), used at 1:500 dilution for immunofluorescence and 1:2000 for western immunoblot; mouse anti-β-actin (Proteintech) used at 1:5000 for western immunoblot. Secondary antibodies were goat anti-mouse IgG (H + L) AlexaFluor-488 (1:1000, A-11001, ThermoFisher Scientific), donkey anti-rabbit conjugated to horseradish peroxidase (1:2000, Cell Signalling Technology) and donkey anti-mouse conjugated to horseradish peroxidase (1:5000, Bio-Rad). Cell nuclei were stained with Hoechst 33342 (1:10,000, ThermoFisher Scientific).

### Chemical Reagents

Metformin for the cell culture experiments was obtained from two sources to counter any batch or manufacturing induced effects (Cambridge Bioscience and APExBIO). For the mouse study the Metformin was prepared from a separate manufacturer (MedChemExpress HY-B0627).

### Immunofluorescent labelling

Cells were washed once with PBS, then fixed in 4% PFA/PBS (Alfa Aesar, Heysham, UK) for minimum of 20 min at room temperature. Cells were then washed three times with PBS and permeabilized in PBST (PBS with 0.2% Triton X-100) for 10 min. Blocking was performed in 2% normal goat serum (GS)/PBST (Dako, Glostrup, Denmark) for 1 h at room temperature. Cells were incubated with primary antibody (diluted in PBST+2% GS) overnight at 4 °C. The following day, cells were washed three times in PBS and incubated with secondary antibodies (diluted in PBST) for 2 h at room temperature in the dark. After three washes with PBS, nuclei were stained in 0.5 μg/mL (1:10,000 Hoechst 33342) for 30 min at room temperature in the dark. Cells were then washed once with PBS and imaged using an Invitrogen EVOS™ M5000 Imaging platform (ThermoFisher Scientific)

### Differentiation assay

Myoblasts were seeded for full confluency at 45,000 cells/well in a black clear bottom 96-well plate (Revvity, Inc) with three technical replicates for each dilution per plate. The next day, growth medium was removed and cells washed with PBS before addition of differentiation medium (DMEM) with Metformin. A serial dilution of Metformin from 5 mM to 50 pM was performed in 1.5ml Eppendorf’s with water. The same volume from each diluent was added to a corresponding 1.5 ml Eppendorf containing the differentiation medium (DMEM) at a ratio of 1:200 to avoid osmotic differences and incubated for the standard differentiation time 72 h.

Cells were imaged on an EVOS and a plate reader. For the plate reader imaging, fluorescence intensities were measured on a ClarioStar microplate reader (BMG Labtech, Ortenberg, Germany) in spectral well averaging scan mode for bottom optics (100 measurements per well, scan diameter 6 mm). Values for the MyHC in each well were divided by the Nuclei to calculate the (MyHC/Nuclei) value for each well. The average of the three technical replicates for each dilution was calculated to be the individual biological replicate value for that dilution.

This plate-reader based assay was used to calculate the expression of myosin heavy chain relative to the number of myonuclei (MyHC/Nuclei) with paired imaging of the self-same well on an EVOS microscope (Fig. 1A). This permits the entire well to be imaged to mitigate imaging area bias.

### EdU proliferation assay

Myoblasts were seeded at proliferation confluency of 5,000 cells/well in a black clear bottom 96-well plate (Revvity, Inc) with three technical replicates for each condition per plate. The following day the medium was removed and growth medium (Promocell) containing either vehicle only (water) or Metformin was added and incubated for 24 h. Myoblasts were then pulsed with EdU reagent (ThermoFisher Scientific) at a 10 μM concentration for 2 h. Myoblasts were then fixed with 4% PFA and the staining protocol undertaken in accordance with the manufacturer’s instructions using the Click-iT^TM^ EdU proliferation Kit (ThermoFisher Scientific), nuclei stained in 0.5 μg/mL (1:10,000 Hoechst 33342) for 30 min at room temperature in the dark, washed once with PBS and imaged using an Invitrogen EVOS™ M5000 Imaging platform (ThermoFisher Scientific). Analysis was performed using StarDist plugin in Fiji to segment the nuclei, counting both total nuclei and nuclei with incorporated EdU and expressed as a percentage of the total.

### Apoptosis, cell Viability/ metabolic activity, ΔΨm, ROS, and MitoROS measurements

45,000 myoblasts were seeded into white, clear-bottom polystyrene 96-well plates, the bottom of the plate covered with a white adhesive seal to enhance luminescence signal intensity (Revvity Inc). The next day, the medium was removed, cells were washed with PBS and then differentiation medium added (DMEM) with Metformin.

Apoptosis was measured by the non-invasive, luminescent RealTime-GloTM Annexin V Assay (JA1001; Promega, Madison, WI, USA). Cell viability/ Metabolic activity was measured using the luminescence RealTime-Glo™ MT Cell Viability assay (Promega). At the 70 h time-point of differentiation, the Assay kit substrates were added to the myotubes for a further 2 hours to the 72 h endpoint. Luminescence intensity measurements were performed on a Mithras LB940 multimode microplate reader (Berthold Technologies, Bad Wildbad, Germany) with either 0.2s or 0.4s exposure. Nuclei were then stained with 0.5 μg/mL (1:10,000 Hoechst 33342) for 30 min. Fluorescent intensity was measured on a ClarioStar microplate reader (BMG Labtech, Ortenberg, Germany) in spectral well averaging scan mode for top optics. The Nuclei values were used to normalize the Annexin V luminesce values.

ΔΨm, ROS, and MitoROS measurements were as previously described^13,17^. At the 72 h endpoint of differentiation: cells were washed twice with 1x Hank’s Balanced Salt Solution (HBSS; with Ca^2+^ and Mg^2+^, Sigma Aldrich), followed by incubation with the ROS/ΔΨm probe for 30 min in the dark at 37°C, 5% CO_2_. General ROS were measured with CM-H2DCFDA (final concentration 2.5 μM, ThermoFisher Scientific), MitoROS with MitoTracker Red CM-H2XRos (final concentration 2.5 μM; M7513, ThermoFisher Scientific), ΔΨm with tetramethylrhodamine methyl ester (TMRM; final concentration 100 nM, ThermoFisher Scientific) and mitochondrial content with MitoTracker Deep Red FM (final concentration 250 nM; M22426, ThermoFisher Scientific). Nuclei were then measured by simultaneous incubation with Hoechst 33342 (final concentration 0.5 μg/mL) for normalisation. After incubation with respective ROS/ΔΨm probes, cells were washed with 1x HBSS, and fluorescence intensities measured on a ClarioStar microplate reader (BMG Labtech, Ortenberg, Germany) in spectral well averaging scan mode (100 measurements per well, scan diameter 6 mm). MitoROS and ΔΨm probe fluorescence was normalised to mitochondrial content (MitoTracker Deep Red fluorescence) after normalisation to input cell quantity, as simultaneously assessed via the respective fluorescence intensities in the same well. ROS probe fluorescence was normalised to input cell quantity. MitoROS and ΔΨm fluorescence measurements are presented as fold change of relative fluorescence intensities after normalisation between controls and samples.

### Protein extraction and western blotting

Myotubes from 6-well plates at 72 h of differentiation were washed in PBS, scraped and pelleted at 300×g, for 3 min and resuspended in cold RIPA buffer (ThermoFisher Scientific) supplemented with a protease and phosphatase inhibitor cocktail (Pierce mini, 1 tab/10 ml; Thermo Fisher Scientific). Cell suspensions were lysed using mechanical shearing by pipetting and vortex mixing and centrifuged (16,000×g for 15 min, 4 °C) to remove nuclei and cellular debris. Protein content was quantified using BCA (Pierce, ThermoFisher Scientific). Protein extracts were separated by SDS-PAGE electrophoresis (pre-cast 4-20% MP TGX Gel, Bio-Rad), transferred to nitrocellulose membrane (0.45 µm, Cytivia) and blocked with 5% milk powder (Sigma Aldrich). Membranes were then divided with a razor blade and immunoblotted with primary antibodies overnight at 4 °C, washed with PBST (PBS + Tween 20) and incubated with species-specific HRP-conjugated secondary antibodies for 2 h at room temperature. HRP activity was detected using Western Lightning Chemiluminescence Reagent (ECL, Cytivia) on a gel imager (ChemiDoc^TM^ MP Imaging system, Bio-Rad).

### Reverse transcription and quantitative PCR (qPCR)

Total RNA was extracted with the RNeasy Mini Kit (QIAGEN) and quantified on a NanoDrop ND-1000 UV/Vis spectrophotometer (ThermoFisher Scientific). For reverse transcription: 1 μg of RNA was used for cDNA synthesis using the QuantiTect Reverse Transcription Kit (QIAGEN) as per manufacturer’s instructions. Quantitative PCR (qPCR) analyses were in triplicate from 5 ng input cDNA on a ViiA7 thermal cycler (Applied Biosystems, Warrington, UK), using Takyon® Low ROX SYBR 2X MasterMix blue dTTP (UF-LPMT-B0701, Eurogentec, Seraing, Belgium) as per manufacturer’s instructions. Target gene expression was normalised to the housekeeper *RPLP0*, and values represented as relative expression (2-ΔΔCT). Presented as fold change after normalisation between controls and samples.

Primer sequences (from Sigma Aldrich): *RPLP0*-fwd: 5’-TGGTCATCCAGCAGGTGTTCGA-3’, *RPLP0*-rev: 5’-ACAGACACTGGCAACATTGCGG-3’, *PINK1*-fwd: 5’-TGACCTTTGCCCCTAACACGAG-3’, *PINK1*-rev: 5’-GTAACTGAACGTGCTGACCCAT-3’, *PARK2*-fwd: 5’-GTGTTTGTCAGGTTCAACTCCA-3’, *PARK2*-rev: 5’-GAAAATCACACGCAACTGGTC-3’, *BNIP3*-fwd: 5’-CGCAGACACCACAAGATACCAA-3’, *BNIP3*-rev: 5’-GCCGACTTGACCAATCCCAT-3’.

### Statistical analysis

Statistical analysis was performed in GraphPad Prism (https://www.graphpad.com/scientific-software/prism/). First, the data were testing for normal distribution (D’Agostino - Pearson normality test). Then, the significance of parametric data was tested by either one-way ANOVA followed by Tukey’s or Dunnett’s post-hoc multiple comparison test, or an unpaired two-tailed t test with Welch’s correction. Experiments were performed at least three independent times (Biological replicates). Detailed statistical information for each experiment, including statistical test, number of independent experiments, *p* values, definition of plotted dots, and error bars are listed in figure legends. For analysis of the force-frequency sigmoidal response curves from the mouse *in vivo* muscle function assay, nonlinear regression was performed using a 4-parameter logistic (4PL) model hill function for the sigmoidal curves. This was followed by curve comparison using an extra sum-of-squares F test, with the additional constrained bottom value to zero for all datasets. This approach evaluated whether datasets were adequately described by a shared curve or required independent parameter estimates. Fitted parameters and their corresponding 95% confidence intervals were extracted from the final model, with *p* value describing the probability of the compared datasets being described by a single response curve.

## Acknowledgements

The authors wish to acknowledge Dr. Ramzi J. Khairallah, co-founder and president of Myologica LLC (https://www.myologica.com) for the performance of the mouse study presented in this work. JG was mainly supported by a grant from Friends of FSH Research awarded to PH and PSZ, with a contribution from UKRI (MR/X001520/1). PH and JQ were supported by UKRI (MR/X001520/1). We are grateful to FSHD Canada (https://fshd.ca) who contracted Myologica to perform the *in vivo* mouse study.

## Author contributions

JG: Conceptualization; Software; Formal analysis; Supervision; Validation; Investigation; Methodology; Project administration; Writing original paper draft, reviewing and editing. JQ: Formal analysis; Investigation. PH: Conceptualization, Funding acquisition. PSZ: Conceptualization, Funding acquisition; Project administration; Writing, reviewing and editing.

## Data availability statement

The data are available from the corresponding author upon reasonable request.

